# Long lasting anxiety following early life stress is dependent on glucocorticoid signaling in zebrafish

**DOI:** 10.1101/2021.05.25.445598

**Authors:** Jacqueline S.R. Chin, Tram-Anh N. Phan, Lydia T. Albert, Alex C. Keene, Erik R. Duboué

## Abstract

Chronic adversity in early childhood is associated with increased anxiety and a propensity for substance abuse later in adulthood, yet the effects of early life stress (ELS) on brain development remains poorly understood. The zebrafish, *Danio* rerio, is a powerful model for studying neurodevelopment and stress. Here, we describe a zebrafish model of ELS and identify a role for glucocorticoid signaling during a critical window in development that leads to long-term changes in brain function. Larval fish subjected to chronic stress in early development exhibited increased anxiety-like behavior and elevated glucocorticoid levels later in life. Increased stress-like behavior was only observed when fish were subjected to ELS within a precise time window in early development, revealing a temporal critical window of sensitivity. Moreover, enhanced anxiety-like behavior only emerges after two months post-ELS, revealing a developmentally specified delay in the effects of ELS. ELS leads to increased levels of baseline cortisol, and resulted in a dysregulation of cortisol receptors, suggesting long-term effects on cortisol signaling. Together, these findings reveal a ‘critical window’ for ELS to affect developmental reprogramming of the glucocorticoid receptor pathway, resulting in chronic elevated stress.

## Introduction

Development and function of the vertebrate brain are influenced by environmental cues and experience in early life (Luby et al., 2020; Reh et al., 2020), yet our understanding of how such environmental cues in specific developmental time windows influences brain development is limited. Chronic stress in early life has robust and long-lasting effects on health and physiology that persist into adulthood (Cowan et al., 2016; Lähdepuro et al., 2019; Taylor, 2010), yet how early life stress (ELS) impacts the developing brain to cause aberrant behaviors in later life remains poorly understood. In mammals, ELS has been shown to cause epigenetic and expression differences in several stress-related genes (Chen et al., 2012; Elliott et al., 2010; McGowan et al., 2009; Murgatroyd et al., 2009; Weaver et al., 2004), and can lead to impaired neuronal proliferation and morphology (Fabricius et al., 2008; Farrell et al., 2016; Korosi et al., 2012; Mirescu et al., 2004; Oomen et al., 2010; Tanapat et al., 1998). These changes impact the function of several brain regions including the hippocampus, amygdala, and hypothalamus, suggesting ELS persistently impacts brain function throughout development (Daun et al., 2020; Elliott et al., 2010; Murgatroyd et al., 2009; Youssef et al., 2019). While its effects are well accepted, a mechanistic understanding of how ELS impairs brain function requires identifying the neuronal changes induced by specific stressors and assessing their impact brain-wide across development.

The zebrafish, *Danio rerio*, is a powerful model for studying how brain development is impacted by stress (Clark et al., 2011; Golla et al., 2020; Hartig et al., 2016). Both behavioral and physiological responses to stress are highly conserved among fish and mammals (Cachat et al., 2010; Canavello et al., 2011). Behaviorally, both adult and larval zebrafish exhibit stereotyped responses following presentation of an aversive or unfamiliar cue including prolonged freezing, reduced exploration, thigmotaxis, and erratic swimming (Maximino et al., 2010a, 2010b). Moreover, several assays have been described and standardized for examining stress in both adults and larvae (Agetsuma et al., 2010; Duboué et al., 2017; Levin et al., 2007; Maximino et al., 2010b). In addition to behavioral reactions to aversive stimuli, fish also display robust physiological responses to stress. Following the presentation of a stressful stimulus, the hypothalamic-pituitary-interrenal (HPI) axis, analogous to the mammalian hypothalamic-pituitary-adrenal (HPA) axis, induces a cascade of events that culminate in the production and release of cortisol (Alsop and Vijayan, 2008; Cachat et al., 2010; Steenbergen et al., 2012; Yeh et al., 2013). Like mammals, cortisol then binds to glucocorticoid and mineralocorticoid receptors in the brain (Alsop and Vijayan, 2008). Combining this fish model of stress with approaches to examine brain development and function has the potential to unravel the mechanistic basis for the effects of ELS on brain development and function.

In this study we induce ELS by applying unpredictable mild electric stimuli at different developmental time points to zebrafish larvae, and measure stress behavior later in juvenile stages. Similar to mammals, ELS in zebrafish at early time points, but not late stages, leads to increased stress behaviors and elevated cortisol levels in later life. Pharmacological analysis of neuroendocrine signaling suggests that ELS disrupts development of cortisol receptors in the brain. Together, these data demonstrate that the effects of ELS are conserved from teleosts to mammals, and point to the zebrafish as a powerful genetic system for studying how ELS impacts brain development, physiology, and function.

## Results

### Zebrafish subjected to ELS have increased anxiety-like behaviors as adults

To determine the long-term consequences of chronic stress in early development on zebrafish, wild-type (AB) (Walker, 1998) larvae were subjected to random pulses of a mild electric current (25V, 200 msec duration, 1 pulse per second) from 2-6 days post fertilization (dpf). This stimulus intensity was chosen as it was the minimum voltage required that caused more than 80% of 2 dpf larvae to react to shock. Moreover, we have previously shown this shock intensity causes a robust stress response in larval zebrafish (Duboué et al., 2017). Control siblings were handled similarly but were not subjected to electric shock. At 6 dpf, following cessation of the ELS protocol, all groups of larvae were transferred to a recirculating system in the main aquatics facility (Figure 1A). The delivery of shock was random, and thus larvae were not able to predict stimulus onset (Figure S1A). Video analysis revealed that larvae react to the electric shocks throughout the 5-day period (Figure S1B). We tested for behavioral differences to stress at larval (7 dpf) and juvenile (60 dpf) stages to determine whether this protocol has a lasting impact on stress response (Figure 1A). Qualitative examination of swim bladder and locomotor behavior under a light microscope revealed that both control and ELS fish appeared healthy immediately following ELS at 7 dpf and after testing behavior at 60 dpf with no gross morphological abnormalities, and swimming in these animals was unaffected (Figure S1C).

**Figure 1.**
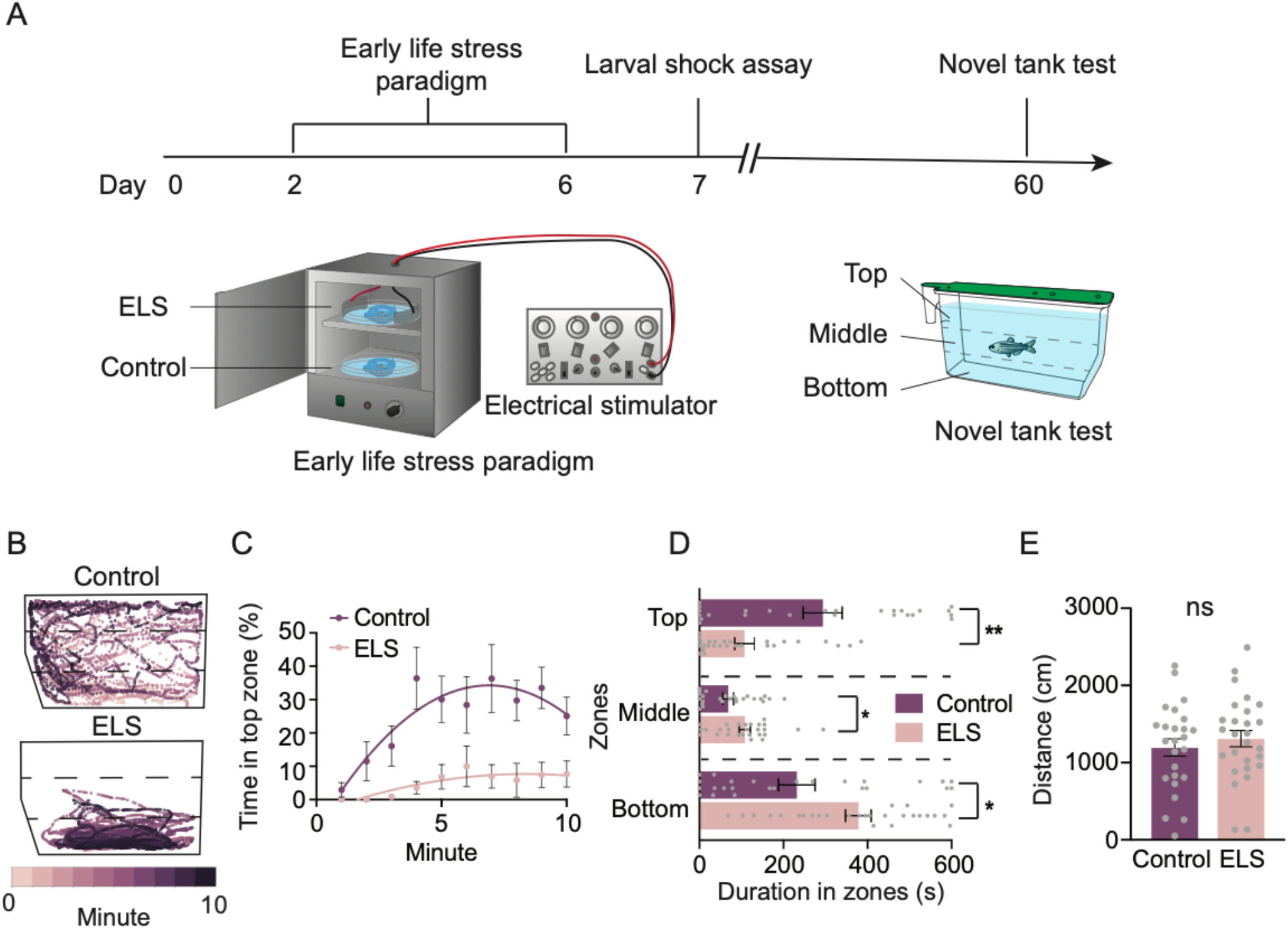
ELS results in exacerbated stress responses in adulthood. (A) Timeline of ELS and behavior experiments and illustrations of the ELS setup and novel tank test. (B) Representative swim trajectories of an individual control and ELS fish revealing that while control fish explored most of the tank, ELS fish swam more at the bottom of the tank.(C) Percentage of time spent in the top zone each minute throughout the 10-min recording. Over time, control animals gradually explored the top zone more frequently than ELS animals. (D) Compared to controls (n= 25), ELS adults (n= 27) spent less time exploring the top (Unpaired t test, p= 0.0003), and more time exploring the bottom (Unpaired t test, p= 0.0038) and middle (Unpaired t test, p= 0.018) zones. (E) Distances travelled were not different between controls (n= 25) and ELS (n= 27) animals (Unpaired t test, p= 0.46). Error bars show ± standard error of the mean. Asterisks denote statistical significance (*: p= 0.05, **: p= 0.005). ns denotes no significance.

To determine if ELS leads to immediate changes in stress behavior, we quantified the differences in the behavioral response to stress between ELS and control animals at 7 dpf, one-day following cessation of shock (Figure S1D). Exposure to mild electric shock causes freezing and reduced locomotion in zebrafish larvae, and the change in freezing and locomotion pre- and post-shock is a reliable measure of stress levels (Duboué et al., 2017). Control animals not subjected to shock in early life displayed significant freezing post-stimulation, revealing a stereotyped stress response in this group at 7 dpf.

Unexpectedly, no significant differences in freezing pre- and post-stimulation at 7 dpf were observed in ELS groups (Figure S1E). As freezing is an indicator of stress, these data suggest that ELS initially causes a reduction in stress responses in larval stages, potentially due to initial habituation. Together, these data suggest that the behavioral effects of ELS may not be immediate, and may emerge only after developmental changes to the brain.

ELS in mammals has been shown to alter stress later in adult stages. To examine whether the effects of ELS are conserved across vertebrates, we measured stress responses in 60 dpf juveniles that were previously subjected to ELS. We applied the novel tank test that is widely used to quantify innate stress behaviors in juvenile and adult fish (Figure 1A) (Cachat et al., 2010; Levin et al., 2007). When introduced into a novel tank, wild-type zebrafish initially prefer the bottom part of the tank, yet over time, they begin to explore the top part with higher frequency; the amount of time spent in the bottom portion of the tank has been validated as a behavioral indicator of stress (Cachat et al., 2010; Facchin et al., 2015; Levin et al., 2007). At 60 dpf, control animals initially prefer the bottom zone of the novel tank, but by 4-min, they spend equal time across all zones (Figure 1B-C). By contrast, adults previously subjected to ELS spent little time in top zone throughout the 10-min recording period (Figure 1B-C). Quantifying total duration spent in all zones revealed that ELS animals spent significantly more time in the bottom zone of the tank over the 10-min recording period, and less time exploring the top zone compared to controls that were not exposed to ELS (Figure 1D). Analysis of total distance moved and total duration immobile revealed no significant differences between control and ELS siblings (Figure 1E & S1F), indicating that bottom dwelling durations were independent of locomotion or lethargy. These data suggest that ELS exposure during early development increases stress behavior in juvenile fish.

### A critical window for increased anxiety following ELS associates with HPI development

Chronic stress in mammals and birds during specified time windows has lasting effects on brain development and function, yet whether this extends to other vertebrates is poorly understood (Elliott et al., 2010; Monaghan et al., 2012; Murgatroyd et al., 2009). To identify whether ELS impacts later stress response through a developmentally sensitive time window, wild-type animals were subjected to the same ELS paradigm described above at varying time periods throughout development (Figure 2A). Fish were subjected to ELS for a 5-day period in the first, second or third week of life (i.e., from 2-6 dpf, 12-16 dpf, or 22-26 dpf) and the effects on stress behavior were measured at 60 dpf (Figure 2A, (a), red box). Whereas chronic stress from 2-6 dpf resulted in enhanced bottom dwelling at 60 dpf, increased bottom-dwelling behavior was not observed when fish were subjected to the ELS paradigm at either 12-16 dpf or 22-26 dpf (Figure 2B; S2A), suggesting wild-type zebrafish are sensitive to ELS in a time window between 2 and 6 days after fertilization.

**Figure 2.**
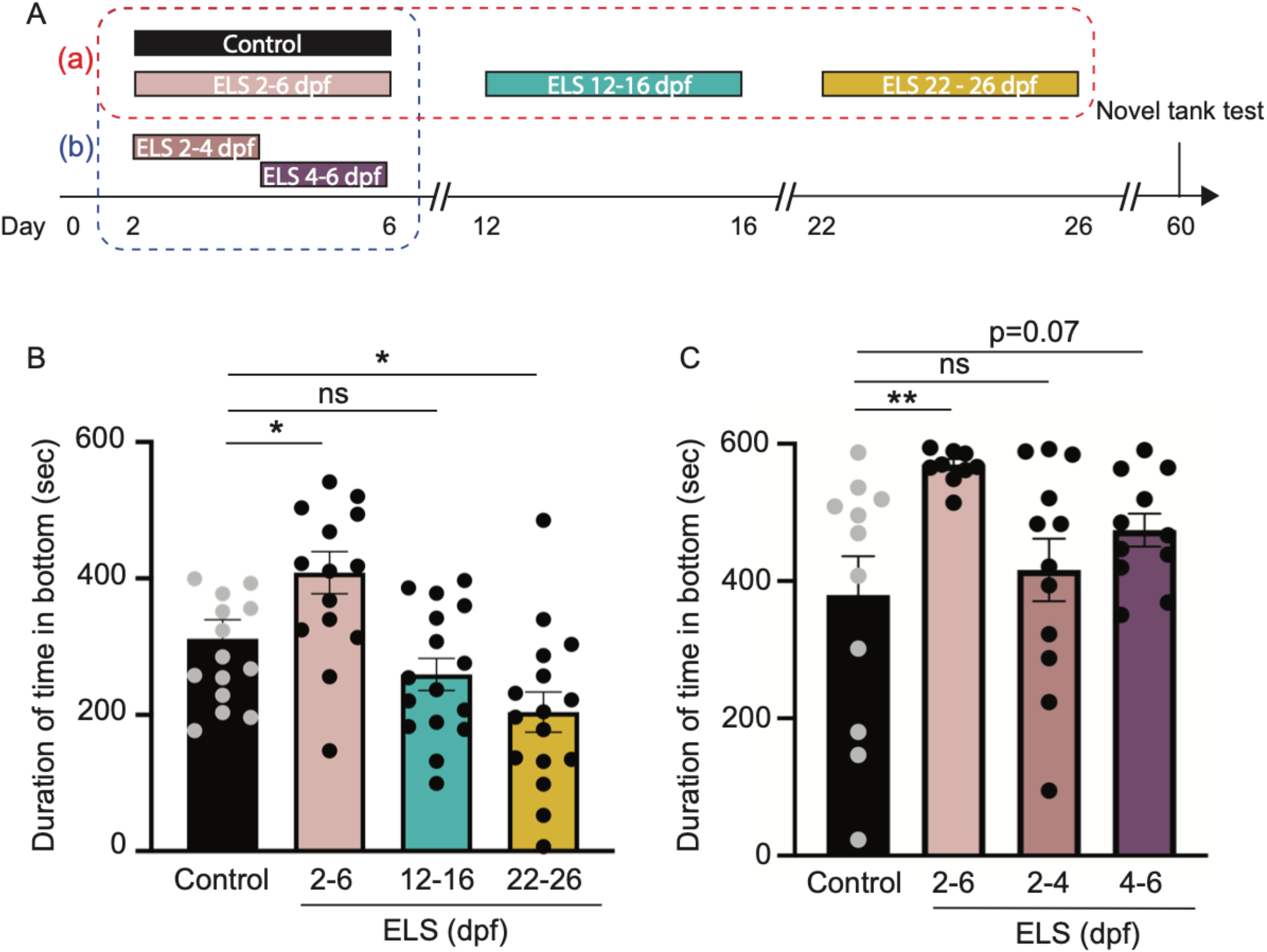
Chronic stress in a time window between 4-6 dpf is critical to impact behavior later in life. (A) Timeline of ELS paradigm and novel tank test. Different groups of larvae were subjected to ELS at different ages. In group (a) (red box), larvae were placed in ELS paradigm at 2-6, 12-16, or 22-26 dpf, alongside same aged and unshocked controls. In group (b) (blue box), larvae were placed in the paradigm at 2-6, 2-4, or 4-6 dpf, while control siblings remained alongside in the incubator throughout the five days. (B) Quantification of duration spent in the bottom zone of the novel tank test at 60 dpf suggest that only chronic stress during the 2-6 dpf period caused increased bottom-dwelling behavior, and not at later times. Multiple unpaired t-tests were performed between control and ELS siblings at three time windows. Control vs. ELS 2-6 dpf : p= 0.014; Control vs. ELS 12-16 dpf: p= 0.081; Control vs. ELS 22-26 dpf : p= 0.0066. (C) Quantification of durations spent in the bottom zone of the novel tank suggest that stress between 4-6 dpf may be sufficient to cause increased bottom-dwelling behavior later in life. Statistical analysis done using multiple unpaired t-tests were performed between control and ELS siblings at three time windows. Control vs. ELS 2-6 dpf: p=0.0026; Control vs. ELS 2-4 dpf: p= 0.31; Control vs. ELS 4-6 dpf: p= 0.07. Controls: n= 11; ELS 2-6 dpf: n= 10; ELS 2-4 dpf: n= 12; ELS 4-6 dpf: n= 11. Error bars show ± standard error of the mean. Asterisks denote statistical significance (***: p= 0.0005, **: p= 0.005, *: p= 0.05).

To define the time window between 2-6 dpf when ELS induces long-term effects on stress response, we subjected larvae to the ELS paradigm described above from 2-6 dpf, or from a more restricted 2-4 or 4-6 dpf time window. Fish were then raised to 60 dpf and tested in the bottom dwelling assay (Figure 2A, (b), blue box). Consistent with previous data, larvae subjected to chronic stress from 2-6 dpf showed significant increases in the total time spent in the bottom of the tank when tested at 60 dpf (Figure 2C). By contrast, no significant effects were observed in zebrafish subjected to shock from 2-4 dpf (Figure 2C). Instead, larvae subjected to chronic shock from 4-6 dpf showed a trend towards increased bottom dwelling behavior when examined at 60 days (Figure 2A, C; Figure S2B, p=0.07). The effect of chronic stress from 4-6 days was less pronounced than when fish were shocked from 2-6; thus, while the main effect is likely in the 4-6 dpf time-period, these data cannot rule out other potentially contributing factors. Together, these data suggest that zebrafish larvae are sensitive to ELS in a critical window after the neuroendocrine stress axis is fully formed and zebrafish are synthesizing their own cortisol, but before brain development is complete (Alsop and Vijayan, 2008, 2009b).

### Increased stress responses are accompanied by increases in basal cortisol levels and expression of stress-related pathway genes

A hallmark of ELS in mammals is prolonged or increased circulating cortisol in the plasma, which correlates with exacerbated stress responses (Goldman-Mellor et al., 2012; Heim et al., 2000; Murgatroyd et al., 2009; Pesonen et al., 2010; Tyrka et al., 2008). We next asked if ELS resulted in changes in the zebrafish neuroendocrine HPI axis. In fish, environmental stressors activate a highly conserved cascade of events from the brain to peripheral tissue to produce and release cortisol (Figure 3A). To determine whether ELS induced alterations in cortisol and other molecules in the HPI axis, we subjected larvae to random shock from 2-6 dpf, raised animals to 60 dpf, and measured both baseline and stress-induced whole-body cortisol levels after being challenged in the novel tank test (Figure 3B). Compared to controls, basal levels of cortisol were significantly increased in ELS animals (Figure 3C). By contrast, no differences were observed between ELS and control animals after the novel tank test (Figure 3C). In both ELS and control siblings, cortisol levels increased significantly following testing in the novel tank test (Figure 3C), suggesting that both groups have an intact physiological response to stress. These data demonstrate that ELS leads to chronically elevated cortisol production.

**Figure 3.**
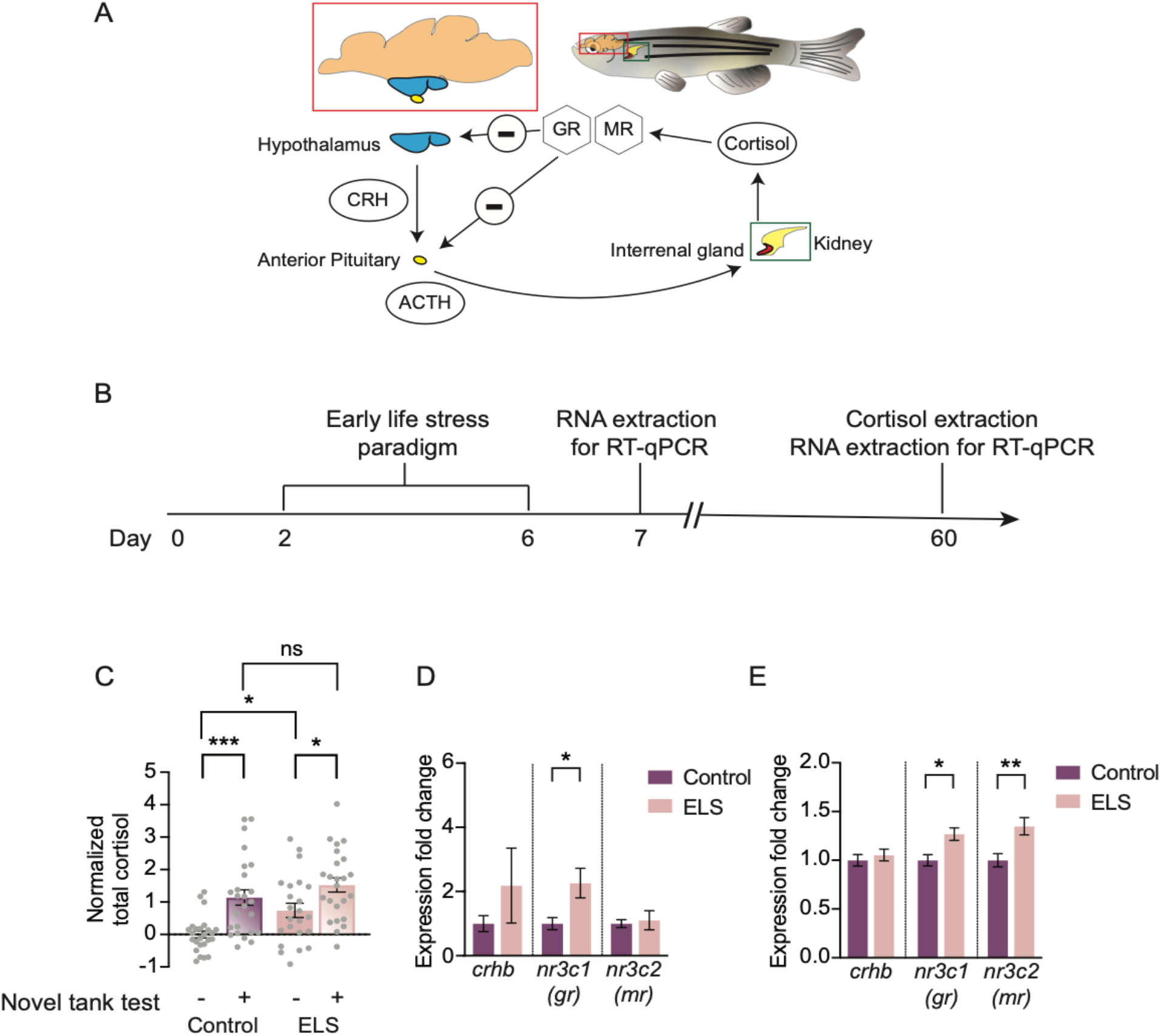
HPI axis is impacted in ELS. (A) The HPI axis, the main stress pathway, and its main genes and effectors. Brain of zebrafish is shown in the red bounding box. Within, the hypothalamus (in blue) and the anterior pituitary (in yellow) is shown. During stress, the hypothalamus signals to the anterior pituitary via CRH, and the anterior pituitary signals to the interrenal gland (shaded in red within the head kidney enclosed in the green bounding box) via ACTH. Cortisol is released from the interrenal gland and binds to GR and MR to negatively regulate its release. (B) Timeline of ELS and experiments performed. (C) Basal cortisol levels (-) were increased in ELS animals (n= 23) compared to controls (n= 25) (One-way ANOVA followed by Sidak’s multiple comparisons post-hoc test, p= 0.044). Elevated cortisol in response to stress (+), after the novel tank test, remain intact in control (n= 25, One-way ANOVA followed by Sidak’s multiple comparisons post-hoc test, p= 0.00040) and ELS adults (n= 24, One-way ANOVA followed by Sidak’s multiple comparisons post-hoc test, p= 0.034). Cortisol levels after stress were no different between controls and ELS animals (One-way ANOVA followed by Sidak’s multiple comparisons post-hoc test, p= 0.54). (D) Quantitative real-time PCR of 7 dpf control (n= 9) and ELS (n= 8) larvae revealed increased gene expression levels of *gr* in ELS animals (Unpaired t test, *gr*: p= 0.018). No significant differences were found in expression levels of *crhb* and *mr* (Unpaired t test, *crhb*: p= 0.31, *mr*: p= 0.74). (E) At 60 dpf, gene expression levels of *gr* and *mr* were increased in brains of ELS animals (Unpaired t test, *gr*: p= 0.0064, *mr*: p= 0.0023), and no difference in *crhb* expression levels (Unpaired t test, p= 0.73) were observed, compared to controls. N= 9 per group. Error bars show ± standard error of the mean. Asterisks denote statistical significance (*: p= 0.05, **: p= 0.005, ***: p= 0.0005). ns denotes no significance.

To further examine how ELS impacts the physiological response to stress, we measured mRNA transcript abundance of several genes in the HPI pathway. Environmental stressors activate *corticotropic releasing hormone* (*crh*) neurons in the hypothalamus (Chandrasekar et al., 2007). CRH then signals indirectly to the interrenal gland to synthesize and release cortisol activates mineralocorticoid (mr) and glucocorticoid receptors (gr) in the brain (Figure 3A) (Bury and Sturm, 2007). To test whether ELS altered the abundance of transcripts in the HPI pathway, larvae were subjected to chronic stress from 2-6 dpf, and then at either 7 or 60 dpf, brains were dissected, and transcript abundance was measured for *crhb, nr3c1* (glucocorticoid receptor; *gr)*, and *nr3c2* (mineralocorticoid receptor; *mr*) using quantitative real-time polymerase chain reaction (qRT-PCR; Figure 3A, B). Quantitative gene expression analysis revealed elevated levels of *gr* in ELS-treated animals compared to control siblings at 7 dpf (Figure 3D), and its expression levels remained significantly higher than controls at 60 dpf (Figure 3E). Furthermore, at 60 dpf, expression levels of *mr* were also elevated in ELS animals (Figure 3E). By contrast, no significant differences were found for the relative expression of *crhb* (Figure 3D, E) suggesting that its broad expression in stress-related regions such as the ventral hypothalamus and preoptic area of the hypothalamus, remains unchanged. Therefore, ELS results in lasting changes in the HPI axis at the transcriptional and physiological levels.

### Enhanced stress following pharmacological activation of the cortisol pathway

In mammals, increased glucocorticoid signaling is strongly correlated with enhanced stress response later in life, yet whether increased glucocorticoids are sufficient to cause enhanced stress in adult stages is unclear. The small size and *ex utero* development of zebrafish has made this system a powerful model for screening of pharmacological compounds (Bruni et al., 2014; Kokel et al., 2010; White et al., 2016). To test whether elevated glucocorticoid signaling alone is sufficient to induce long-term changes in stress response, we pharmacologically activated glucocorticoid signaling between 2-6 dpf, and measured whether this was sufficient to cause increased anxiety later in juvenile fish. Glucocorticoids were provided in a continuous flow system that provided larvae boluses of drug that mimicked the electric shock protocol used to induces ELS (Figure 4A).

**Figure 4.**
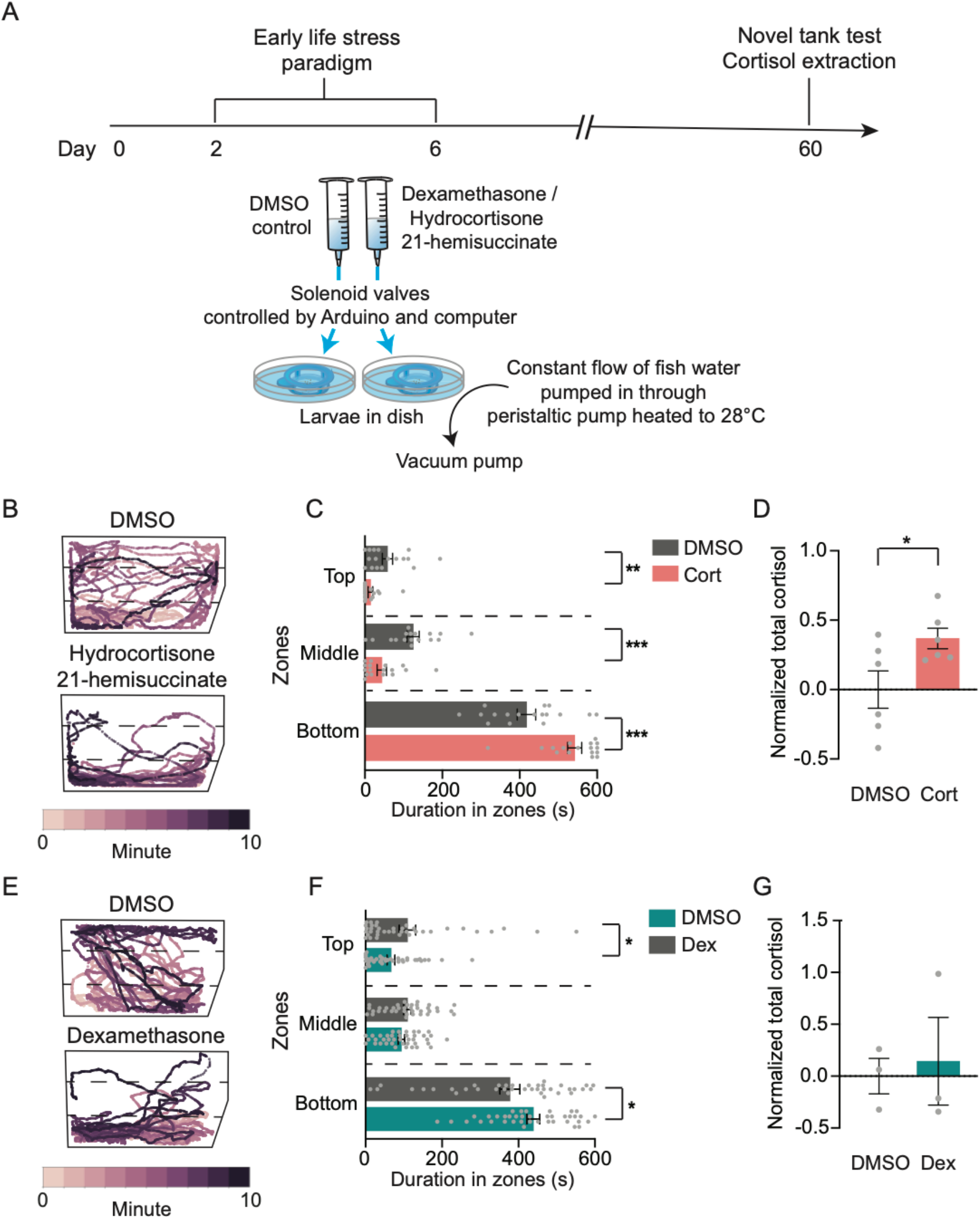
Treatment with corticosteroid receptor agonists in early life induces adulthood anxiety. (A) Timeline of drug-treated ELS and experiments conducted. A schematic diagram of the setup is presented below the timeline. (B) Representative swim paths of 60 dpf controls (DMSO) and hydrocortisone 21-hemisuccinate-(Cort) treated animals. Drug-treated individuals tend to spend more time at the bottom of the tank. (C) Total duration spent in the bottom zone of the novel tank was significantly increased in Cort animals (Unpaired t test, p= 0.0001), and decreased in the top (Unpaired t test, p= 0.0026) and middle (Unpaired t test, p= 0.0002) zones, compared to control DMSO. N= 17 per group. (D) Basal cortisol levels were significantly higher in Cort animals than control DMSO siblings (n= 6 per group, Unpaired t test, p= 0.038). (E) Representative swim paths of 60 dpf controls (DMSO) and dexamethasone-(Dex) treated animals. Drug-treated individuals tend to spend more time at the bottom of the tank. (F) Quantification of durations spent in top, middle, and bottom zones of the novel tank test revealed that Dex animals spent more time in the bottom zones (Unpaired t test, p= 0.025), and less time in the top (p= 0.035) than control DMSO animals yet no difference was observed in time spent in the middle (Unpaired t test, p= 0.11). DMSO: n= 38; Dex: n= 40. (G) Measurements of basal cortisol levels were no different between Dex and DMSO animals (n= 3 per group, Unpaired t test, p= 0.77). Error bars show ± standard error of the mean. Asterisks denote statistical significance (*: p= 0.05, **: p= 0.005, ***: p= 0.0005). ns denotes no significance.

Juvenile animals dosed with the synthetic glucocorticoid that binds to both MR and GR hydrocortisone 21-hemisuccinate (Cort) in early life spent increased time in the bottom and less time in the top of the novel tank compared to undosed sibling controls (Figure 4B & C). Likewise, we observed an increase in basal levels of cortisol in Cort-treated animals compared to controls (Figure 4D). Locomotor activity was reduced in Cort-treated animals (Figure S4A), but this is unlikely to be attributed to a general loss of coordination because the total time spent immobile did not differ from control siblings (Figure S4B).

Taken together, these findings reveal that pharmacological activation of the MR/GR pathways phenocopy shock-induced ELS.

We next asked if overactivation of GR alone was sufficient to phenocopy ELS through the application of dexamethasone (Dex), a selective glucocorticoid agonist (Axelrod, 1976; Brookes et al., 2012). Dex-treated animals spent more time in the bottom of the novel tank and less time in the top and middle zones compared to control animals (Figure 4E & F). Interestingly, basal cortisol levels were not significantly different between Dex-treated and control animals (Figure 4D), suggesting cortisol levels may be separable from the stress response. Distance travelled and duration of immobility did not differ between control DMSO- and Dex-treated animals (Figure S4C & D). Taken together, these data suggest that ELS alters brain development through dysregulation of GR signaling.

## Discussion

Our findings demonstrate that zebrafish, like mammals, have augmented stress responses when subjected to ELS early in development. We also show that the negative effects of ELS act, in part, through chronic activation of GR, whose expression changes over the course of development (Alsop and Vijayan, 2008). Notably, enhanced stress following ELS was not observed in the day following ELS, but rather enhanced stress emerged later, further supporting the notion that ELS is impacting development of the brain. The critical window for chronic stress is after the time where the neuroendocrine stress axis is formed (Alsop and Vijayan, 2009a), suggesting that ELS is not acting directly on the development of the HPI axis, but rather on hormonal signaling to the brain.

In humans, childhood trauma, such as abuse or parental neglect, has potent impacts on adult behavior, stress responsivity, susceptibility to develop stress-related disorders, and substance abuse issues (Chapman et al., 2004; Fogelman and Canli, 2019; Gluckman et al., 2008), and genome-wide association studies have identified components of glucocorticoid-mediated gene regulatory networks suggesting ELS may alter the expression of this pathway during brain development. Genetic manipulation of the glucocorticoid receptor in early life, but not later life, also leads to enhanced stress-related pathologies in later life, supporting the notion that glucocorticoid receptor signaling in early development is a critical mediator of ELS (Ridder et al., 2005; Wei et al., 2004). However, an analysis of how abnormal glucocorticoid signaling alters the development of the brain has been challenging in mammalian systems. Our findings reveal the molecular basis of ELS is highly conserved and provide a powerful genetic model for investigating the mechanistic basis of ELS on neurodevelopment.

### A novel model of ELS

Here, we demonstrate that zebrafish display significantly elevated stress responses when subjected to chronic stressors in larval stages, similar to what is found in mammals. This suggest zebrafish may be a unique and powerful complement to mammalian systems, especially for studying how ELS causes changes to neurodevelopment. Zebrafish have several attributes that make them an attractive complement to mammalian models in the study of ELS. Originally established as a model in development, zebrafish fertilization and development are ex-utero and fish embryos are transparent (de Abreu et al., 2020; Patton and Zon, 2001). The transparency not only permits examination of development ex-utero, but also highlights the power of the system in circuit neuroscience (Friedrich et al., 2010, 2013). This work sets the stage for using powerful tools, such as whole-brain functional imaging and genetic manipulation of precise neuronal subsets, to study how brain anatomy and function change over the course of development in response to ELS (Ahrens and Engert, 2015; Ahrens et al., 2012; Muto et al., 2011).

### The role of GR in enhanced stress following ELS

Several hypotheses exist to explain how enhanced cortisol activation can impair development. A prevailing model suggests that the effects of ELS emerge through an imbalance in the relative proportion of MR:GR (Kitraki et al., 2004; de Kloet et al., 1998, 2018). Our data do not support this hypothesis in zebrafish. While the expression of both GR and MR were higher in ELS-subjected animals, the relative balance, or ratio, was not significantly different (Figure S3). Alternatively, overactivation of GR alone may impair brain development. In mammals, chronic activation of GR results in neuronal death, dendritic spine retraction, and reduced spike frequencies (Joëls and de Kloet, 1991; Joels et al., 1991; Macleod et al., 2003; McCullers and Herman, 1998; Sapolsky et al., 1988; Woolley et al., 1990, 1991). Moreover, overexpression of GR throughout the life of the rodents, or transient activation of GR only in early development is sufficient to lead to enhanced anxiety, whereas transient activation in adult stages has no effect (Wei et al., 2004). Our data thus support a conserved role of dysregulated GR signaling in enhanced risk, and point to an ancient origin for GR signaling in the brain and its contribution to stress disorders.

### Neurodevelopmental implications of ELS in zebrafish

A central strength of the zebrafish in the study of ELS is the ability to interrogate the effects of stressors at specific developmental time points, and associate those time points that impact stress to precise developmental processes. This central strength reveals several exciting findings about ELS. First, our data reveal that long lasting effects of chronic stress emerge only when stressors are given in early time points. Thus, enhanced stress following ELS is likely not a passive response to elevated stress but rather ELS likely impedes normal brain development. Moreover, that larvae are particularly sensitive to stressors from 4-6 dpf, and less so at 2-4 dpf, points to specific developmental processes that may be impacted.

The HPI axis begins forming early in development, and expression levels of both GR and MR show significant fluctuations until approximately 2 dpf; by 49 hours post fertilization, expression levels of GR in normal reared animals are stable. Zebrafish do not begin to produce cortisol in response to exogenous stressors until 4 dpf (Alsop and Vijayan, 2008). Because the HPI axis is fully formed by 4 dpf, our data suggest that ELS is not impacting the development of the neuroendocrine stress axis. Moreover, as larvae are particularly sensitive to ELS from 4-6dpf, a time point when animals are beginning to produce cortisol, it is likely that ELS is leading to overactivation of glucocorticoid signaling, which in turn impacts brain development.

Significant changes in brain development also occur during the 4-6 dpf window. In the zebrafish forebrain, a large neuroanatomical region with loci analogous to the mammalian limbic system, newborn neurons begin to form by 2 dpf (Mueller et al., 2011). By 4 dpf, development of the zebrafish forebrain is complete and neuronal properties such as spontaneous activity and neurotransmitter identity are beginning to develop (Heisenberg et al., 1996). Significant change in neuronal activity also occur in these early time points. Spontaneous activity in the zebrafish brain is observed beginning from 2 dpf through adulthood, yet significant changes emerge over developmental time. In the tectum, spontaneous activity is random and disorganized at 2 dpf, yet by 8 dpf, the circuit is functional mature, and clusters of functionally relevant neurons fire in unison (Boulanger-Weill and Sumbre, 2019; Thomas Pietri and Romano, 2017). These data support the notion that development and function of the neuroendocrine stress axis are impacted by ELS. Future studies utilizing transgenic and neuroimaging techniques on this model of ELS will reveal how specific changes in brain development and maturation of neuronal activity following ELS lead to enhanced stress responses in later life.

## Conclusions

Our data introduce a new model in the field of early life stress, and uncover several principles about how ELS may impact the developing brain. The zebrafish is a strong model in developmental biology and neuroscience, and thus the unique strengths of this model provide unprecedented insight into how the brain responds to stress in these early time periods. Zebrafish exhibit stereotyped and well-studied stress responses and there are a number of assays available to examine stress in fish. Moreover, the high conservation at the neuronal and physiological levels between fish and humans suggest that findings in zebrafish will translate well to the mammalian system, and should complement mammalian work. Furthermore, the large collection of mutant and transgenic lines available in zebrafish, optical approaches for monitoring brain development in vivo, computational tools such as the recently developed brain atlases, and whole-brain functional imaging all coalesce and provide a single model that uniquely bridges development and neuroscience.

---

**Figure S1.**
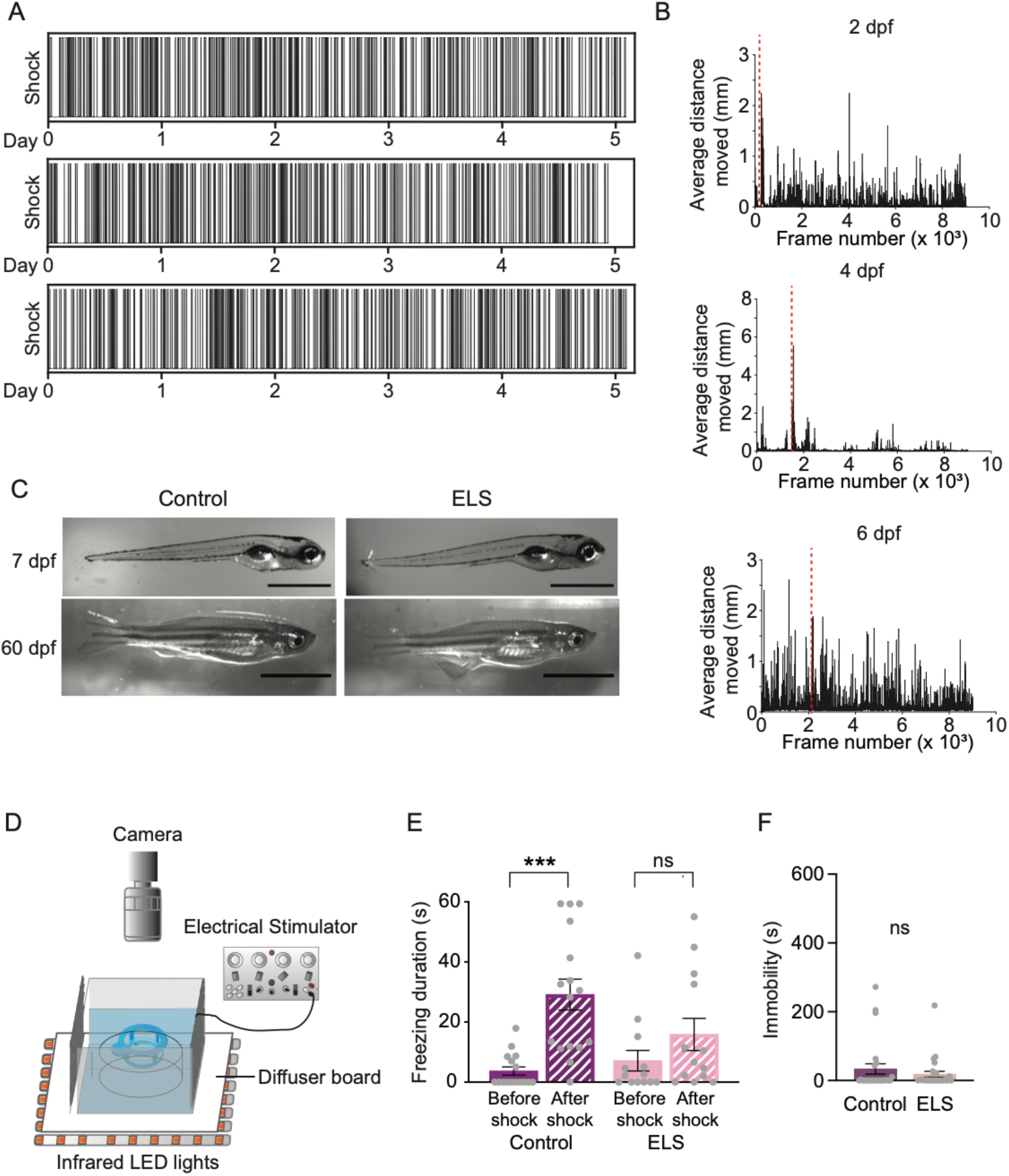
Mild electric shocks in ELS paradigm are random and cause large movements in larvae upon stimulation; related to Figure 1. (A) Representative graphs of random mild electric shocks applied throughout the five-day setup using a custom written script. Each line in the graphs represent an instance of shock applied. (B) Average distance moved in a ten-minute recording of 2, 4, and 6 dpf larvae in the ELS paradigm where a period of electric shocks (red dashed lines) were given. Electric shocks were accompanied by increase in distance moved in the frames following the shocks. 6 dpf larvae on average moved more throughout the recording, even when no shock was given. N= 16 per group. (C) Images of control and ELS siblings at 7 and 60 dpf revealed no gross physical defects. Scale bars in 7 and 60 dpf images represent 1mm and 5mm, respectively. (D) Diagram of shock assay performed on 7dpf larvae, a day after the ELS paradigm. Locomotor behavior was recorded a minute before shock stimuli and a minute immediately after the stimuli. (E) 7 dpf ELS larvae showed dampened ‘freezing’ response to electric shock compared to control larvae. Statistical analysis was done using the Kruskal Wallis test followed by Dunn’s multiple comparisons post-hoc test. Controls: n= 16; ELS: n= 13. Control before shock vs. Control after shock: p=0.0002; ELS before shock vs. ELS after shock: p= 0.84; Control before shock vs. ELS before shock: p> 0.99; Control after shock vs. ELS after shock: p= 0.19. (F) Quantification of total duration of immobility in the novel tank test at 60 dpf showed no difference between ELS (n= 27) and control (n= 25) siblings (Unpaired t test, p= 0.36). Error bars show ± standard error of the mean. Asterisks denote statistical significance (***: p= 0.0005). ns denotes no significance.

**Figure S2.**
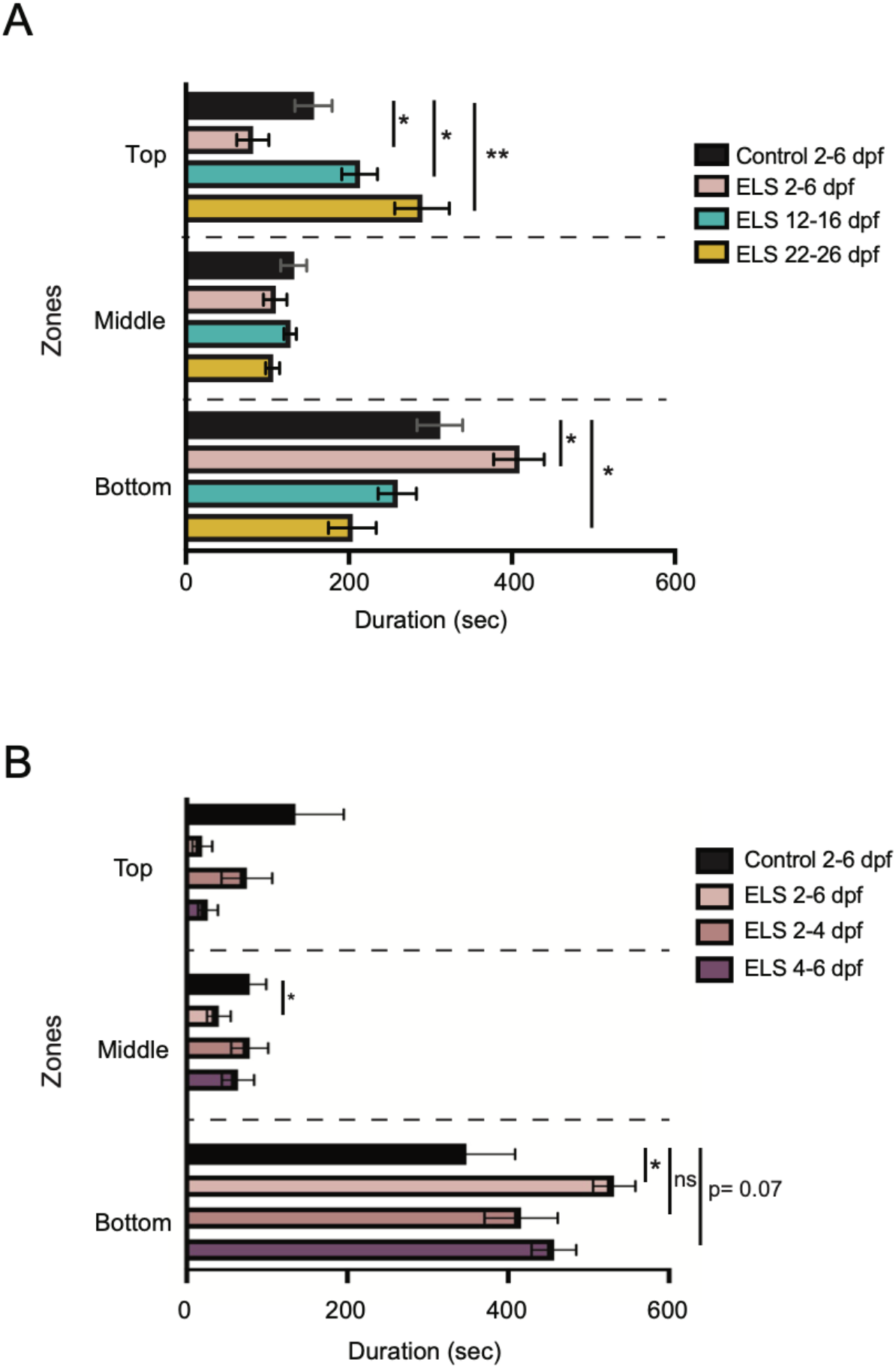
Quantification of durations in all zones in a novel tank test; related to Figure 2. (A) Quantification of durations in all zones revealed that only chronic stress in early life impacts anxiety behavior late in life. Multiple unpaired t-tests were performed between control and ELS siblings at three time windows. Top: Control vs. ELS 2-6 dpf: p=0.0098; Control vs. ELS 12-16 dpf: p= 0.041; Control vs. ELS 22-26 dpf: p= 0.0015; Middle: Control vs. ELS 2-6 dpf : p= 0.014; Control vs. ELS 12-16 dpf: p= 0.40; Control vs. ELS 22-26 dpf : p= 0.078; Bottom: Control vs. ELS 2-6 dpf : p= 0.014; Control dpf vs. ELS 12-16 dpf: p= 0.081; Control dpf vs. ELS 22-26 dpf : p= 0.081. Control: n= 15; ELS 2-6 dpf: n= 15; ELS 12-16 dpf: n= 16; ELS 22-26 dpf: n= 16. (B) Quantification of durations spent in the three zones in the novel tank suggested that stress between 4-6 dpf may be sufficient to cause increased bottom-dwelling behavior later in life. Multiple unpaired t-tests were used to compare between groups in each zone. Top zone-Control vs. ELS 2-6 dpf: p=0.041; Control vs. ELS 2-4 dpf: p= 0.37; Control vs. ELS 4-6 dpf: p= 0.065. Middle zone-Control vs. ELS 2-6 dpf: p=0.0056; Control vs. ELS 2-4 dpf: p= 0.43; Control vs. ELS 4-6 dpf: p= 0.13. Bottom zone-Control vs. ELS 2-6 dpf: p=0.0026; Control vs. ELS 2-4 dpf: p= 0.31; Control vs. ELS 4-6 dpf: p= 0.07. Controls: n= 11; ELS 2-6 dpf: n= 10; ELS 2-4 dpf: n= 12; ELS 4-6 dpf: n= 11. Error bars show ± standard error of the mean. Asterisks denote statistical significance (*: p= 0.05, **: p= 0.005). ns denotes no significance.

**Figure S3.**
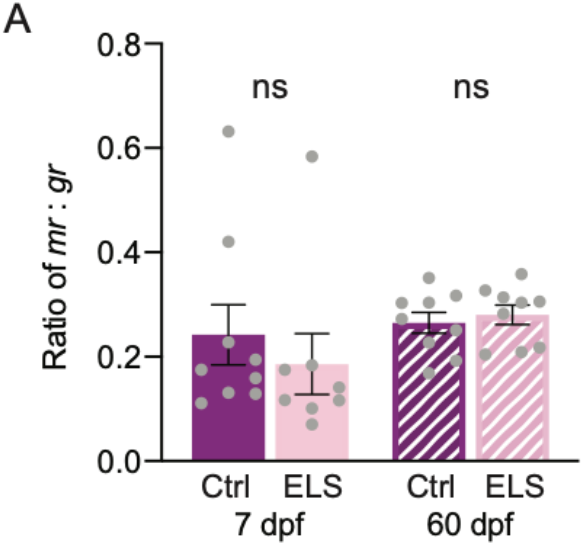
Analysis of MR: GR ratio at 7 and 60 dpf revealed no differences between controls and ELS animals; related to Figure 3. Statistical analysis was done using the Mann-Whitney test. 7dpf - Control: n= 9; ELS: n= 8, p= 0.20. 60 dpf – Control: n= 9; ELS: n= 9, p= 0.49. ns denotes no significance.

**Figure S4.**
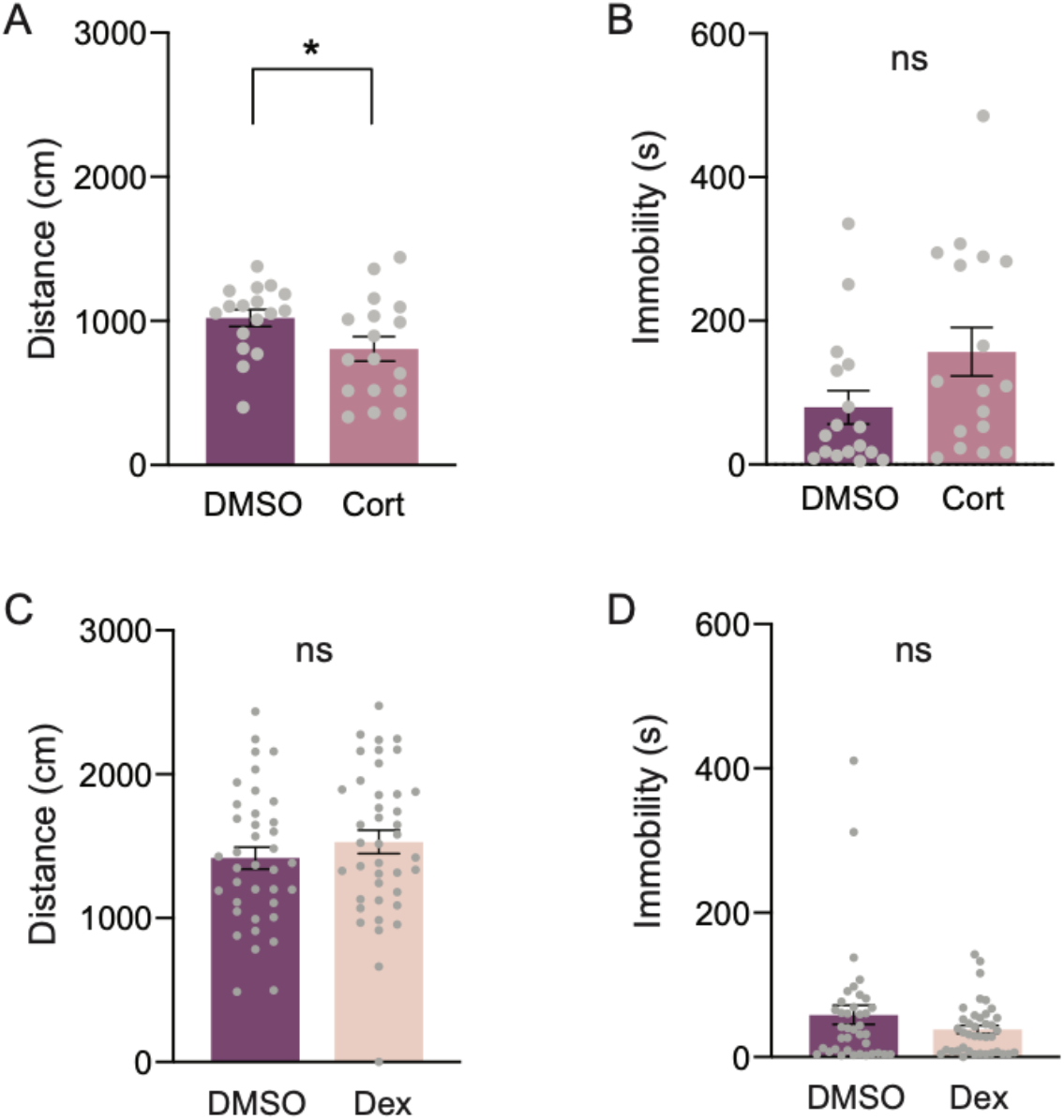
Analysis of locomotor behavior of drug-treated animals; related to Figure 4. (A) Total distance traveled was significantly reduced in Cort individuals compared to control DMSO siblings (Unpaired t test, p= 0.045). N= 17 per group. (B) Total durations spent immobile were no different between Cort and DMSO siblings (Unpaired t test, p= 0.068). N= 17 per group. (C) Analysis of distance traveled in the novel tank revealed no significant differences between control DMSO (n= 38) and Dex (n= 40) animals (Unpaired t test, p= 0.32). (D) Similarly, control DMSO (n= 38) and Dex (n= 40) siblings spent comparable amounts of time being immobile (Unpaired t test, p= 0.16). Asterisks denote statistical significance (*: p= 0.05). ns denotes no significance.

**Table S1.**
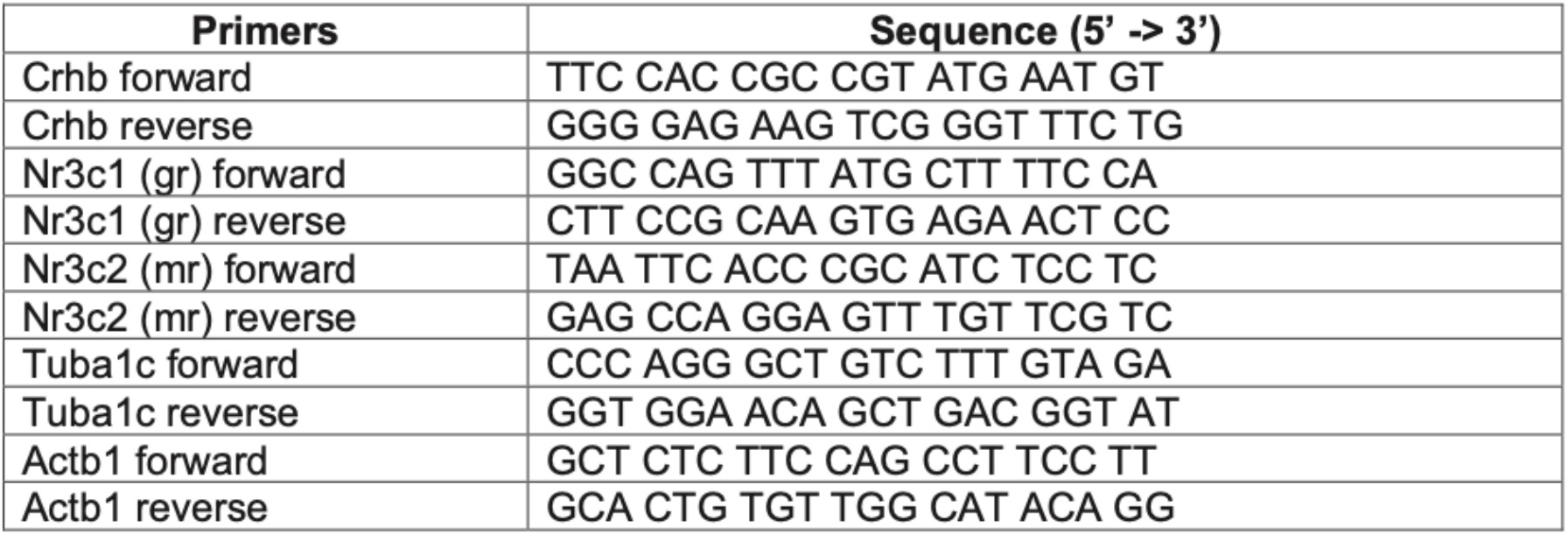
List of RT-qPCR primers; related to Figure 3.

## STAR★ METHODS

### Resource Availability

#### Lead Contact

Further information and requests for resources and reagents should be directed to and will be fulfilled by the Lead Contact, Erik R. Duboué.

#### Materials Availability

This study did not generate new unique reagents.

#### Data and Code Availability

The datasets and code supporting the current study have not been deposited in a public repository, but are available from the corresponding author on request.

### Experimental Model and Subject Details

Wild-type AB (Walker, 1998) zebrafish were used in all experiments in this study. Adults were kept in 1.8-10 L tanks on a recirculating aquatics system (Aquaneering). The water temperature was maintained at 28 ± 1 °C, and the light cycle was set to a 14:10 light:dark cycle. In order to breed larvae, 4-6 adult breeder fish (half males and half females) were placed in a mating tank (1L, ZHCT100, Aquaneering), together with some plastic plants, and allowed to spawn naturally overnight. The morning after spawning, eggs were collected, placed in 100 × 15mm Petri dishes (Fisherbrand, Fisher Scientific Inc.), and housed in an incubator (Heratherm IMC18, Thermo Scientific) set at 28 ± 1 °C. Fish subjected to ELS, as well as their control siblings, were housed on the same re-circulating system, in dedicated 1.8 L tanks containing 500 mL of 5 ppt salt water, co-cultured with L-type rotifers from 6 to 15 dpf and fed with GEMMA Micro 75 (Skretting Zebrafish) daily. At 15 dpf, the valve was opened to allow system water to flow at a slow rate into the tanks, rotifers were washed out, and fish transitioned to be fed *Artemia* (brine shrimp) and GEMMA Micro 150 (Skretting Zebrafish). Fish older than 30 dpf were fed GEMMA Micro 300 (Skretting Zebrafish) daily. Experiments were carried out between 11 a.m. to 6 p.m., and according to a protocol (A17-22) approved by the Institutional Animal Care and Use Committee of Florida Atlantic University.

### Method Details

#### ELS paradigm

Early life stress was induced in zebrafish larvae using a custom written computer program (MATLAB R2019a, MathWorks). This script controlled a square pulse stimulator (SD9 Grass Stimulator, Grass Technologies Inc) interfaced through a microcontroller board (Arduino Uno R3, Arduino) to randomly deliver electric current to fish from 2-6 dpf. Groups of thirty to fifty 2 dpf larvae were placed in 40 µm cell strainers (Corning Inc.) placed in 100 × 15 mm Petri dishes (Fisherbrand, Fisher Scientific Inc.) filled with fresh system water. A dish containing control larvae, which were not subjected to ELS, as well as a dish containing larvae that were subjected to ELS were placed in an incubator at 28°C on a 14:10 light:dark cycle. A pair of electrodes connected to the square pulse stimulator was placed on opposite ends of the dish containing larvae to be subjected to ELS. The stimulator was controlled by a microcontroller board (Arduino Uno R3, Arduino), which was connected to a computer (Inspiron 15 3000 series, Dell). The custom written program was designed to evaluate a random number between 0 and 1 every ten minutes. In cases where the random number was greater than or equal to 0.5, 5-electric pulses were delivered (25 V, 1Hz, 200 msec pulse duration), effectively delivering shocks 50% of the time; by contrast, if the random number was less than 0.5, no stimulation was provided. The net effect of this program was random stimulation so that fish were not able to predict. Fish were subjected to this protocol from 2-6 dpf. At the end of the 6 dpf, fish were removed from the incubator, and placed on the recirculating system where they were raised to 60 dpf. For critical time windows experiments, fish at 2 dpf, 12 dpf, and 22 dpf were subjected to the ELS paradigm as described above for five days, except that for 12 and 22 dpf fish, twenty juveniles were placed in glass Pyrex bowls (470mL, Pyrex) and co-cultured with L-type rotifers and fed with GEMMA Micro 75 (Skretting Zebrafish) from 12-16dpf, and GEMMA Micro 150 (Skretting Zebrafish) from 22-26 dpf.

#### Pharmacological induction of ELS

ELS was induced pharmacologically using a modification of the ELS paradigm described above. Thirty 2 dpf larvae were placed in 40 µm cell strainers (Corning Inc.) housed in in petri dishes (50 × 15 mm, Fisherbrand, Fisher Scientific Inc.). Petri dishes were placed in an enclosed space, and were connected to a flowthrough, temperature-controlled water delivery system, consisting of a peristaltic pump (EW-78001-60, Cole-Parmer), which delivered fresh system water at a rate of 30 µL/sec, a pair of heating/cooling cubes (ALA Scientific Instruments), that maintained water temperature at 28 ± 1 °C, and a vacuum pump (ALA Scientific Instruments), which maintained water volume of 12 ± 1 mL in the dishes. Using the same computer program and microcontroller board previously described, two solenoid valves connected to dosing syringes were programmed to deliver either 1.5 mL of 0.1mM dexamethasone (PHR1526, Sigma-Aldrich), 25µM hydrocortisone 21-hemisuccinate (H4881, Sigma-Aldrich), or control 1% dimethyl sulphoxide (D8418, Sigma-Aldrich) into each petri dish. The treatments were delivered at 50% chance at every 10 min interval. Similarly, at the end of 6 dpf, the larvae were placed back into the facility and raised to 60 dpf.

#### Larval shock assay

At 7 dpf, a day after the ELS paradigm, the shock assay was performed as previously described (Duboue et al) to analyze stress behavior. Briefly, a single larva was placed in a 40 µm cell strainer (Corning Inc.) that was on a 3cm raised platform in a 6 × 6 × 6 cm (L x W x D) container. On opposite sides, two electrodes were connected to a stimulator (SD9 Grass Stimulator, Grass Technologies Inc.). The bottom of the container was illuminated by infrared Light Emitting Diodes (LED; 880 nm) and a white acrylic board that acted as a diffuser. Overhead, a high framerate cMOS camera (Grasshopper3, PointGrey, FLIR Integrated Imaging Systems, Inc.) captured larval behavior at 120fps for 2.1 min - a minute of normal swimming behavior, followed by five seconds of electric shock at 1 pulse-per-second at 25V, and then a minute post-shock. Records of each trial were captured using FlyCap2 Software (version 2.11, PointGrey, FLIR Integrated Imaging Systems, Inc.). Tracking of individuals were done offline in EthoVision XT (v13, Noldus), and each frame was inspected manually for inaccurate tracking. Raw XY coordinates were exported and a custom-written script in MATLAB (R2019a, MathWorks) was used to analyze ‘freezing’ behavior. Freezing was determined as immobility for more than 1.99 secs, as previously described (Duboue et al., 2017). Larvae used for this behavioral assay were euthanized and not used in adult tests and analyses.

#### Novel tank test

60 dpf (± 3 days) control and experimental adults were transported in their home tanks from the fish facility to the behavior room and acclimated for an hour before the behavior test. An adult was first subjected to a stressor of being removed out of water for 3 min with a net. Then, the individual was allowed to recover briefly for 10 min in a 250 mL beaker filled with 200 mL fresh system water. Following the rest period, the individual was gently poured into a 1.8L novel tank (ZT180, Aquaneering) positioned with custom designed infrared Light Emitting Diodes lights (LED; 880 nm) in the background, and recorded for 10-min in front of a high-frame-rate cMOS camera (Grasshopper3, PointGrey, FLIR Integrated Imaging Systems, Inc.). Records of each trial were captured using FlyCap2 Software (version 2.11, PointGrey, FLIR Integrated Imaging Systems, Inc.). Tracking of individuals and analyses of durations spent in zones were done offline in EthoVision XT (v13, Noldus), and each frame was inspected manually for inaccurate tracking. The height of the water level was divided equally into three zones to analyze the durations spent in each zone. Durations spent in each zone, distance travelled, duration of immobility, and XY coordinates were obtained from the software. From XY coordinates obtained, swim trajectories were plotted using matplotlib in Python (v3.7).

#### Cortisol measurements

To obtain basal cortisol, 60 dpf adults were flash frozen in individual 1.5 mL Eppendorf tubes before the novel tank test. Post-stress cortisol was obtained from adults immediately after the novel tank test. Frozen samples were kept at -20°C for up to a month before cortisol extractions. Whole adult fish were weighed, then homogenized, and cortisol was extracted following Cachat et al., 2010 with minor modifications. Briefly, each individual was homogenized in 500 µL phosphate buffered saline (PBS). Then, another 500 µL PBS was added before decanting the sample into a glass scintillation vial. To extract cortisol, 2 mL diethyl ether (E134, Fisher Scientific) was added to the homogenate. Next, the samples were vortexed and then centrifuged. Subsequently, the organic diethyl ether layer containing cortisol was extracted into a new glass vial. The extraction steps were performed three times. After evaporation of the organic layer, 1 mL PBS was added and the sample was kept at 4°C overnight. The ELISA assay (#1-3002, Salivary Cortisol ELISA Kit, Salimetrics, LLC) measuring the amounts of cortisol was performed the following day. A standard dilution curve was made with standards provided. Total cortisol was normalized to body weight. Then, to account for daily variations in extractions and variations in kits, cortisol levels were normalized again to the mean control basal cortisol levels for each day using the following formula:

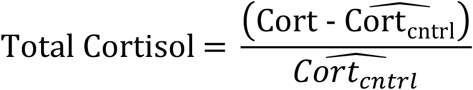

where *Cort* represents total baseline cortisol levels, and 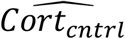 indicates the average cortisol levels measured for each day.

#### Quantitative gene expression analysis of HPI axis genes

For gene expression analysis at 60 dpf, brains were dissected and immediately snap frozen in liquid nitrogen. Three brains were pooled together for each biological replicate, and were kept in -80°C for RNA extraction the following day. RNA was extracted using TRIzol (Thermo Fisher Scientific) and the RNeasy mini kit (QIAGEN). Genomic DNA was removed by DNase treatment (RNase-free DNase set, QIAGEN). 1000μg of RNA was reverse transcribed into cDNA (iScript cDNA Synthesis Kit, Bio-Rad), and the subsequent cDNA was diluted to a concentration of 50 ng/μL to use for quantitative real-time PCR (CFX96 Touch Real-Time PCR Detection System, Bio-Rad). For gene expression analysis at 7 dpf, groups of twenty larvae were used for each biological replicate. Target gene expression levels were normalized to actin beta-1 and tubulin alpha-1c. Refer to Table S1 for primer sequences.

### Quantification and Statistical Analysis

Statistical analyses were performed using Prism 8 (v8.0.2, GraphPad Software). Parametric tests were used unless the data failed the Shapiro-Wilk normality test, then non-parametric tests were used. For pairwise comparisons between control and ELS groups, one-tailed unpaired t tests were used. The non-parametric equivalent, Mann-Whitney test was used when the data failed the normality test. Where comparisons were made between multiple groups, one-way ANOVA was performed, and when statistical significance between groups was obtained, the Sidak’s multiple comparisons post-hoc test was performed. The non-parametric equivalent of the one-way ANOVA used was the Kruskal-Wallis test, followed by the Dunn’s multiple comparisons post-hoc test.

### Key Resources Table

| REAGENT or RESOURCE | SOURCE | IDENTIFIER |
| --- | --- | --- |
| Chemicals, Peptides, and Recombinant Proteins |  |  |
| Diethyl ether | Fisher Scientific | Cat#E134 |
| Dexamethasone | Sigma-Aldrich | Cat#PHR1526 |
| Hydrocortisone-21 hemisuccinate sodium salt | Sigma-Aldrich | Cat#H4881 |
| Dimethyl sulphoxide | Sigma-Aldrich | Cat#D8418 |
| Critical Commercial Assays |  |  |
| Human salivary cortisol ELISA kit | Salimetrics | Cat#1-3002 |
| Experimental Models: Organisms/Strains |  |  |
| AB |  | RRID: ZIRC_ZL1 |
| Oligonucleotides |  |  |
| RT-qPCR primers for *crhb*, see Table S1. | This paper | N/A |
| RT-qPCR primers for *gr*, see Table S1. | This paper | N/A |
| RT-qPCR primers for *mr*, see Table S1. | This paper | N/A |
| RT-qPCR primers for *tubulin alpha-1c*, see Table S1. | This paper | N/A |
| RT-qPCR primers for *actin beta-1*, see Table S1. | This paper | N/A |
| Software and Algorithms |  |  |
| Python 3.7 (graphs of swim paths and Figure S1C) | Python | https://www.python.org/ |
| Prism 8 (graphs and statistics) | GraphPad | https://www.graphpad.com/ |
| Ethovision XT (offline tracking of individuals) | Noldus | https://www.noldus.com/ethovision-xt |
| MATLAB R2019a (ELS paradigm script) | MathWorks | https://www.mathworks.com/products/matlab.html |
| Other |  |  |
| Arduino Uno R3 | Arduino | Model# A000066 |
| Solenoid Valve | RobotGeek | Item# # HW-WVALVE |

